# Comparative genomics reveals the origins and diversity of arthropod immune systems

**DOI:** 10.1101/010942

**Authors:** William J. Palmer, Francis M. Jiggins

## Abstract

While the innate immune system of insects is well-studied, comparatively little is known about how other arthropods defend themselves against infection. We have characterised key immune components in the genomes of five chelicerates, a myriapod and a crustacean. We found clear traces of an ancient origin of innate immunity, with some arthropods havingToll-like receptors and C_3_-complement factors that are more closely related in sequence or structure to vertebrates than other arthropods. Across the arthropods some components of the immune system, like the Toll signalling pathway, are highly conserved. However, there is also remarkable diversity. The chelicerates apparently lack the Imd signalling pathway and BGRPs – a key class of pathogen recognition receptors. Many genes have large copy number variation across species, and this may sometimes be accompanied by changes in function. For example, peptidoglycan recognition proteins (PGRPs) have frequently lost their catalytic activity and switch between secreted and intracellular forms. There has been extensive duplication of the cellular immune receptor Dscam in several species, which may be an alternative way to generate the high diversity that produced by alternative splicing in insects. Our results provide a detailed analysis of the immune systems of several important groups of animals and lay the foundations for functional work on these groups.

## Introduction

All animals must defend themselves against a battery of natural enemies, ranging from pathogens such as viruses, bacteria and fungi, to macroscopic parasites such as parasitic worms or insects. The immune defences that have evolved in response to this challenge must distinguish self from non-self, and produce effectors that target and kill these invaders. All the major groups of animals possess an innate immune system, where immune receptors are genetically hard-coded and the response is typically relatively non-specific with respect to individual pathogen strains or previous exposure (Hoffmann et al. 1999; Kimbrell and Beutler 2001). The innate immune system originated early in animal evolution before the split between protostomes and deuterostomes, as some components of the vertebrate innate immune system show clear homology to insect immune molecules(Hoffmann and Reichhart 2002; Wang et al. 2014; Zhu et al. 2005). In addition to an innate immune system, jawed vertebrates possess an adaptive or acquired immune system, where receptor diversity is generated somatically and there is immunological memory(Hoffmann et al. 1999; Kimbrell and Beutler 2001; Janeway and Medzhitov 2002; Hoffmann and Reichhart 2002; Smith et al. 2011). This would seem to be an evolutionary novelty, as key components of the adaptive immune system are not found much beyond jawed vertebrates(Söderhäll 2011).

Arthropods have a powerful innate immune response, our understanding of which comes largely from insects, especially *Drosophila* and mosquitoes. In these species, pathogen associated molecular patterns (PAMPs)(Janeway and Medzhitov 2002) such as bacterial peptidoglycan or fungal beta-1,3 glucan are recognised by pattern recognition receptors like peptidoglycan binding proteins (PGRPs) and beta-1,3 glucan binding proteins (BGRPs)(Kimbrell and Beutler 2001; Lemaitre and Hoffmann 2007; Waterhouse et al. 2007). Following recognition, these receptors then activate the Toll and Imd signalling pathways, leading to the translocation of Nf-kb transcription factors into the nucleus and a humoral response characterised by the expression of antimicrobial peptides(Lemaitre and Hoffmann 2007). In addition there is a melanisation response that kills parasites by depositing the dark pigment melanin along with the production of toxic molecules (Lemaitre and Hoffmann 2007). Alongside the humoral response, there is a cellular responses involving phagocytosis and the encapsulation of larger parasites in layers of blood cells(Lemaitre and Hoffmann 2007).

Whole-genome analyses have revealed much conservation of key immune pathways and gene families between insect species. The Toll, Imd, JAK/STAT and JNK signalling pathways are remarkably well conserved, often in 1:1 orthologous relationships between species(Tanaka et al. 2008; Gerardo et al. 2010; Zou et al. 2007; Evans et al. 2006; Waterhouse et al. 2007). A notable exception to this pattern is the pea aphid, which appears to have lost the Imd pathway(Gerardo et al. 2010). Despite this, much variation in presence/absence, copy number and sequence divergence is observed in other genes, particularly those encoding recognition and effector molecules(Gerardo et al. 2010; Waterhouse et al. 2007; Sackton et al. 2007). For example, mosquitoes show extensive duplications in gene families associated with the response to the malaria parasite *Plasmodium*(Waterhouse et al. 2007).

Beyond the insects, Toll-like receptors (TLRs), their associated signalling components, and Nf-kb transcription factors all have mammalian homologues, suggesting the origin of these genes predates the protostome/deuterostome split over six hundred million years ago(Janeway and Medzhitov 2002; Hoffmann and Reichhart 2002; Hoffmann et al. 1999; Lemaitre and Hoffmann 2007). The same is true for the PGRPs and thioester-containing proteins (TEPs), which show similarities to vertebrate alpha-2 macroglobulins and complement factors(Sekiguchi, Fujito, and Nonaka 2012; Zhu et al. 2005). Components of the Imd pathway resemble the tumour necrosis factor receptor (TNFR) pathway of mammals(Hoffmann 2003).

The evolution and diversity of innate immune systems across the arthropods remains poorly understood, despite the importance of arthropods as disease vectors, pests, and components of biodiversity. As yet, the only detailed whole-genome analysis of a non-insect arthropod investigated the crustacean *Daphnia,* which is the sister group to the insects(McTaggart et al. 2009). This found a repertoire of immune genes that is remarkably insect-like, with the notable absence of peptidoglycan recognition proteins. However, there is a lack of genome-level studies of the more divergent myriapods and chelicerates(Gerardo et al. 2010; Grbić et al. 2011). As such the timing and nature of many key innovations in arthropods remains unresolved, and we lack an overview of the immune system in major arthropods groups.

The recent sequencing of multiple whole arthropod genomes, some of which are unpublished, provides an opportunity to examine the arthropod immunity gene repertoire in a systematic and consistent fashion. In this study we have used these genomes to characterise the evolution of the innate immune system across all the main arthropod taxa. Our results show both remarkable diversification of the immune response across the arthropods, and unexpected conservation and similarities to mammalian genes.

## Results and Discussion

To investigate the evolution and origins of the arthropod innate immune system, we identified homologs of insect immunity genes in species that diverged early in the evolution of the arthropods. The arthropods are a phylum that contains four extant sub-phyla, with the chelicerates, the myriapods and the crustaceans sequentially diverging from the lineage leading to the insects. This allows us to identify which components of the immune system were present in the ancestral arthropod, and which have been gained or lost later in evolution.

The timing of events early in the evolution of the arthropods is highly uncertain, but it is clear that these four major groups all diverged very early in the evolution of animals. The common ancestor of the arthropods existed an estimated 543 million years ago (ma), and by 511 ma all four of the subphyla had formed(Rota-Stabelli, Daley, and Pisani 2013); see figures below). In our analysis we included the genomes of the insect *Drosophila melanogaster*, the crustacean *Daphnia pulex* (Water Flea), the myriapod *Strigamia maritima* (Coastal Centipede) and five species of chelicerate: *Mesobuthus martensii* (Chinese Scorpion)(Cao et al. 2013)*, Parasteatoda tepidariorum* (House Spider), *Ixodes scapularis* (Deer Tick), *Metaseiulus occidentalis* (Western Orchard Predatory Mite), and *Tetranychus urticae* (Red Spider Mite)(Grbić et al. 2011).

To identify homologs of immunity genes across these great phylogenetic distances, we combined methods based on sequence similarity with the predicted cellular location of proteins and analyses of domains, motifs and residues that are known to be essential for the immune function of the encoded proteins (Table 1). These features can often be identified across very distantly related species, which allows us to guard against the inevitable loss of power to detect sequence similarity when looking at distantly related species.

### Arthropod Toll-like receptors are a dynamically evolving gene family that includes relatives of vertebrate TLRs

Toll-like receptors (TLRs) are transmembrane proteins that play a central role in the immune response of insects and vertebrates. They have an extracellular leucine-rich repeat (LRR) region at the N-terminal, and a cytosolic Toll/interleukin-1 receptor (TIR) domain at the C-terminal (Leulier and Lemaitre 2008). In mammals different TLRs directly recognise a variety of pathogen-associated molecular patterns. In *Drosophila* Toll-1 plays key roles in both development and immunity, with the immune function relying on Toll-1 binding to an endogenous cytokine rather than directly to pathogen molecules (see below; (Leulier and Lemaitre 2008)). Several other *Drosophila* TLRs have been suggested to have immune functions, although these are poorly characterised, and it is likely that most have primarily developmental roles(Leulier and Lemaitre 2008).

**Figure 1.**
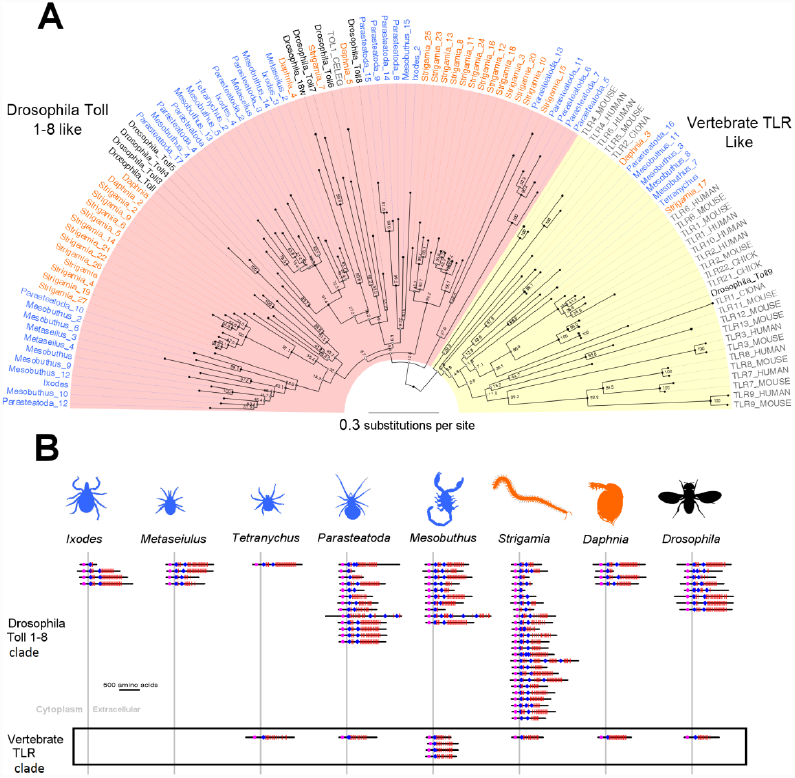
Arthropod Toll-like receptors (TLRs). (A) Phylogenetic tree of TLRs from seven species of arthropods, four chordates (human, mouse, chicken and *Ciona*), and the nematode *Caenorhabditis*. The tree was reconstructed by maximum likelihood from the TIR domains and is midpoint rooted. Myriapod and crustacean taxon labels are orange, chelicerates blue and *Drosophila* black. Node labels are bootstrap support. (B) The domain structure of arthropod TLRs. TLRs in the yellow vertebrate clade of panel A are shown in the black box. Red bars are leucine-rich repeats, blue diamonds are cysteine clusters, magenta the TIR domain, and the grey line represents the plasma membrane. Domain locations are all to scale.

In all the arthropod genomes we find multiple TLRs, with a TIR domain separated from LRRs by a transmembrane helix. On a tree reconstructed from the sequence of the TIR domain, Toll-1 clusters with three other *Drosophila* TLRs (albeit with weak support; Figure 1), so there are no clear 1:1 homologs of Toll-1 outside of the insects. Therefore, it cannot be predicted which if any of the TLRs beyond the insects are likely to have a role in immune signalling.

The TLRs have been frequently lost and duplicated during arthropod evolution, as there is extensive copy number variation and little congruence between the gene and species trees (Figure 1). Within the myriapods there have been two large copy number expansions, resulting in a total of 27 TLRs in the *Strigamia* genome (Figure 1). The spider *Parasteatoda* and scorpion *Mesobuthus* also have a high number of TLRs (16 and 14 respectively), while at the other extreme the tick *Ixodes* has just two (Figure 1).

There are two major structural classes of TLRs and both are widespread in arthropods. The sccTLRs have a single cysteine cluster the end of the LRRs adjacent to the cell membrane, while the mccTLRs have multiple cysteine clusters(Imler and Zheng 2004; Leulier and Lemaitre 2008). The vertebrate TLRs are largely sccTLRs, while *Drosophila* Toll receptors other than Toll 9 are mccTLRs(Imler and Zheng 2004; Leulier and Lemaitre 2008). We found that while mccTLRs are most common in arthropods, vertebrate-like sccTLRs are also widespread (Figure 1).

The division between sccTLRs and mccTLRs is reflected in their evolutionary relationships, with the two structural classes forming two major clades on the TLR phylogeny (Figure 1). In most cases the arthropod sccTLRs are more closely related to vertebrate sccTLRs than they are to arthropod mccTLRs. (Figure 1). The mccTLR clade (figure 1, pink shading, ‘Drosophila Toll 1-8 like’) contains eight of the nine *Drosophila* Tolls, along with most of the arthropod TLRs that we identified and the nematode TLR *Caenorhabitis* TOL1 (nematodes are protostomes like arthropods). The sccTLR clade (figure 1, yellow shading, ‘Vertebrate TLR like’) contains all sequences from the four chordate deuterostomes we included in this analysis — human, mouse, chicken and ciona(Sasaki et al. 2009) — but also sccTLRs from *Drosophila, Daphnia* and *Strigamia* and several chelicerate sequences. This pattern of the structure of the TLRs being reflected in their phylogeny is especially striking as the phylogeny is reconstructed using the sequence of the intracellular TIR domain, while the structural classification is based on extracellular sequences. The structural classification also provides strong corroboration for this phylogenetic division, despite the tree being poorly resolved (Figure 1). Therefore, throughout the arthropods there are a small number of TLRs that are more similar to vertebrate TLRs than other arthropod TLRs.

The presence of arthropod TLRs clustering with vertebrates shows that mccTLRs and sccTLRs diverged very early in animal evolution, but it is unclear whether the common ancestor of protostomes and deuterostomes had both types of TLR. Our tree places arthropod sccTLRs sequences interspersed among vertebrate TLRs (Figure 1). However, this may simply reflect error in the tree reconstruction, as bootstrap support for these relationships is low and we were unable to reject a tree where all the deuterostome and protostome taxa were monophyletic (Shimodaira-Hasegawa Test: 2∆*l*=6, *p*=>0.05). If our midpoint root to the tree is correct (Figure 1), then the common ancestor of deuterostomes and protostomes had both classes of TLR. However, if it is not, then the common ancestor may have had just sccTLRs, with mccTLRs appearing later in protostome evolution.

Studies of the immune function the *Drosophila* sccTLR (Toll 9) have produced conflicting results, so it remains uncertain what the function of these vertebrate-like TLRs is in arthropods(Narbonne-Reveau, Charroux, and Royet 2011; Ooi et al. 2002). We also find a few cases of unusual extracellular domain structures. A small number of TLRs in the mccTLR clade that only have a single cysteine cluster adjacent to the plasma membrane. In all but one case these are short truncated proteins. There are also a small number of TLRs with more than two cysteine clusters.

### The Toll signalling pathway is conserved across arthropods

The humoral immune response of *Drosophila* and other insects centres on the Toll and Imd pathways, both of which result in Nf-kb transcription factors being activated and translocated into the nucleus, where they upregulate the expression of antimicrobial peptides and other genes. In *Drosophila,* recognition of Gram-positive bacteria and fungi by BGRPs and their short-chain interacting PGRPs cause cleavage of spatzle, which subsequently binds to and activates Toll-1, initiating the Toll pathway(Bosco-Drayon et al. 2012; Gendrin et al. 2013)(Bosco-Drayon et al. 2012; Gendrin et al. 2013)^7,28^.

Following binding by cleaved spatzle, the Toll-1 TIR domain initiates a signalling cascade that culminates in the translocation of the Nf-kb transcription factors Dif and dorsal to the nucleus(Lemaitre and Hoffmann 2007).

**Figure 2.**
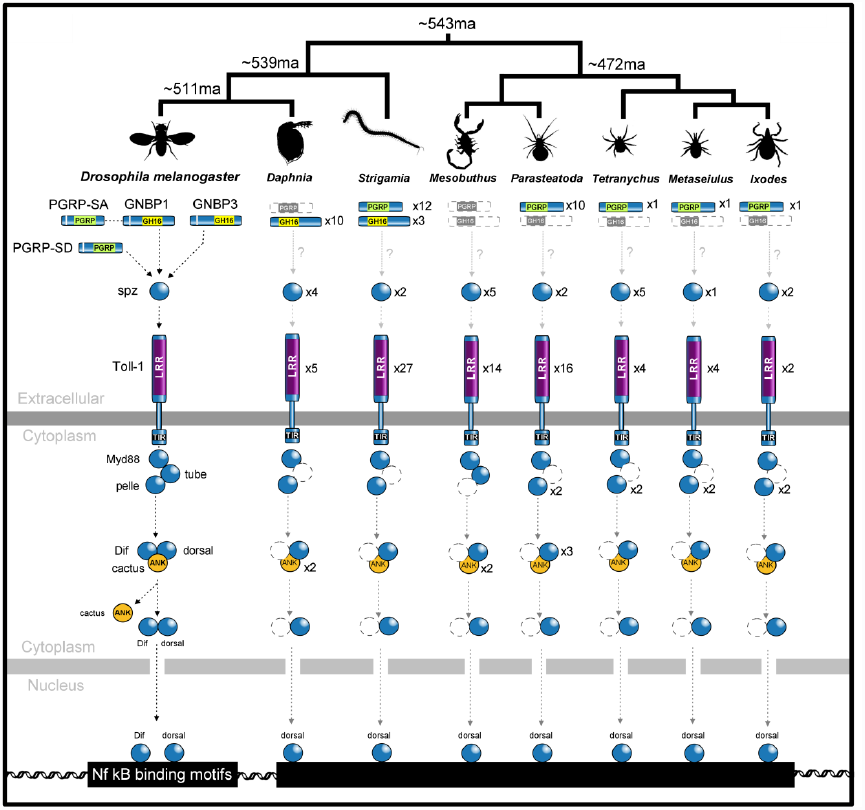
Presence or absence of Toll pathway members across the arthropods. Dashed grey symbols represent proteins where no homolog was detected. Divergence dates on the phylogeny are in millions of years before present (ma) and from **(Rota-Stabelli, Daley, and Pisani 2013)**. Key domains are shown (Ank: ankyrin repeat, LRR: leucine rich repeat)

We found Toll pathway members to be highly conserved across the arthropods (Figure 2), with homologues of spatzle, Myd88, pelle, cactus and dorsal in all species. We failed to find a Tube homologue in any of the species studied except *Mesobuthus*, but this is likely to be a lack of power to detect the gene, as a previous analysis of Tube suggested that it is homologous to IRAK-4, which occupies an equivalent position in the vertebrate Toll pathway(Towb, Sun, and Wasserman 2009). Indeed we find multiple multiple proteins across species with the IRAK-like death domains that characterise Tube. Most genes are present in single copies, although the Nf-kb transcription factor Dorsal has been duplicated twice in the spider *Parasteatoda* (as well as being duplicated in *Drosophila*), and its inhibitor Cactus is duplicated in *Daphnia* and *Mesobuthus.*

### The Imd signalling pathway is highly reduced in chelicerates

In *Drosophila* the humoral immune response to Gram-negative bacteria is controlled by the Imd pathway, which is initiated by the binding of the transmembrane protein PGRP-LC to peptidoglycan. The intracellular RHIM motif of PGRP-LC interacts with Imd, initiating the signalling cascade(Meister et al. 2009). Imd in turn activates TAK1, which together with the IKKb/y complex, Fadd and DREDD (a caspase-8 homologue) activate the Nf-kb transcription factor Relish. Relish is translocated into the nucleus, up-regulating antimicrobial peptides and other genes. Imd also activates the JNK pathway through Tak1(Hoffmann 2003; Lemaitre and Hoffmann 2007; Waterhouse et al. 2007).

*Relish* is an unusual and easily identifiable gene, as the Nf-kB transcription factor and its inhibitor are combined in a single protein(Wang et al. 2014). We find clear Relish homologs in all but one species, suggesting that a Relish-based immune response may have been present in the common ancestor of the arthropods (Figure 3). The IkB kinase complex (IKK) is required for the cleavage and activation of Relish(Hoffmann 2003; Lemaitre and Hoffmann 2007; Waterhouse et al. 2007), and we homologs of both the catalytic subunit, IKKb, and the regulatory subunit, IKKg, in most species (Figure 3). The Relish homologs all have an N-terminal relish-like domain that shows closest sequence similarity to *Drosophila* Relish, however some lack the distinctive C-terminal ankyrin repeat region that plays the role of the Nf-kB inhibitor (Figure 3). In *Daphnia* the ankyrin repeat region is absent from the public gene model due to an error in the automated gene model prediction, as manual annotation identified an additional portion of the gene with ankyrin repeats(McTaggart et al. 2009).It is unclear whether the three chelicerate sequences lacking the ankryin repeats are also annotation errors or reflect true loss of this region.

Relish appears to have been lost entirely in the mite *Metaseiulus* (Figure 3), where no homologues could be discerned by sequence similarity or conserved domain searches (RHD-n relish domain). This is especially striking as *Metaseiulus* has a small and well-assembled genome(Hoy 2009) so this is unlikely to be an artefact of an incomplete genome sequence. Overall, this species is missing more Imd pathway components than any of the other species (Figure 3), supporting the hypothesis that this branch of the immune response may have been lost. This would not be a unique occurrence, as Relish and other Imd pathway components have also been lost in the pea aphid(Gerardo et al. 2010).

**Figure 3.**
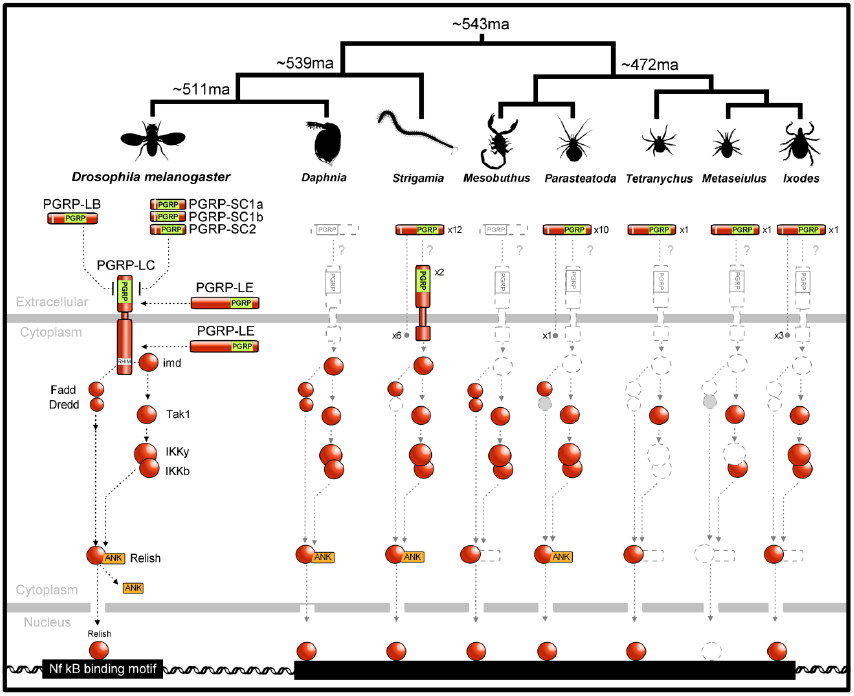
Presence or absence of Imd pathway members across the arthropods. Dashed grey symbols represent genes where no homolog was detected. Divergence date in millions of years before present (ma) are from (Rota-Stabelli, Daley, and Pisani 2013). ANK represents ankyrin repeats.

Despite the conservation of Relish in most species, many other key components of the Imd pathway were only found in the mandibulates and were absent from the chelicerates (Figure 3). In the mites and ticks (*Tetranychus, Metaseiulus* and *Ixodes*), we fail to find any likely Imd, Fadd or Dredd homologues. In arachnids (*Parasteatoda* and *Mesobuthus*) we find possible Fadd and Dredd homologues, although they apparently lack N-terminal DED domains, so they may not have the same function as in *Drosophila*. As Fadd homologues are known to be widespread in the animal kingdom, these results suggests that secondary losses of Fadd and Dredd may have occurred in the chelicerates.

The absence of an intact Imd pathway in the chelicerates is supported by the distribution of transmembrane PGRPs, which are found at the start of the pathway in *Drosophila.* The PGRP domain can clearly be detected across these phylogenetic distances (see below), and transmembrane proteins can be robustly predicted. Transmembrane PGRPs were entirely absent from the chelicerates (Figures 3 and 4).

Although the Imd pathway is largely intact in both the myriapods and crustaceans, transmembrane PGRPs are not. *Daphnia* does not possess any PGRPs (Figures 3 and 4; (McTaggart et al. 2009)), suggesting that the Imd pathway is either not functional or is activated in a different way in this species. In the myriapod *Strigamia* there are two transmembrane PGRPs, but we failed to identify the Imd-interacting RHIM domain using HMMER. Therefore, the activation of the Imd pathway by a transmembrane PGRP could either represent an innovation acquired in the insect lineage, or it may have been lost in the crustaceans.

In contrast to the absence of many Imd pathway components in chelicerates, the JNK pathway is highly conserved across the arthropods with Basket and Jun universally present. This may be a consequence of this pathway playing a role in many key cellular processes in addition to its role in Imd-related signalling.

### The JAK/STAT signalling pathway is highly conserved

The JAK/STAT pathway plays a role in the immune response of both mammals and *Drosophila.* In *Drosophila* and *Anopheles* mosquitoes, following immune challenge activated STAT translocates to the nucleus where it alters the expression of many genes, including upregulating the *Drosophila* immunity protein TEP1(Agaisse and Perrimon 2004). We find clear homologues of Domeless, Hop, and Stat92e in most species (Supplementary Table 3). *Parasteatoda* and *Mesobuthus* and *Metaseiulus* lack a clear Hop homologue however and Domeless would also appear to be absent from *Mesobuthus*.

### Peptidoglycan-recognition proteins (PGRPs)

PGRPs bind to bacterial peptidoglycan and can act as pathogen recognition receptors, negative regulators of the immune response, or effectors that kill bacteria(Gendrin et al. 2013; Bosco-Drayon et al. 2012; Lemaitre and Hoffmann 2007). They fall into two main groups. The non-catalytic PGRPs can function as pattern recognition receptors, and play a key role in activating the Toll and Imd pathways of *Drosophila* following infection (PGRP-LC/LE in the Imd pathway and PGRP-SA/SD in the Toll pathway). The catalytic PGRPs possess amidase activity that allows them to enzymatically break down peptidoglycan (in *Drosophila:* PGRP-SC1/2, SB1/2, and LB), and can they function either as negative regulators of the immune response by removing immunogenic peptidoglycan or as effectors that kill bacteria by degrading their peptidoglycan bacteria(Gendrin et al. 2013; Bosco-Drayon et al. 2012; Lemaitre and Hoffmann 2007). PGRPs are also found in mammals, and were therefore presumably present in the common ancestor of the arthropods(Bischoff et al. 2006; Zaidman-Rémy et al. 2006; Lemaitre and Hoffmann 2007; Zaidman-Rémy et al. 2011).

**Figure 4.**
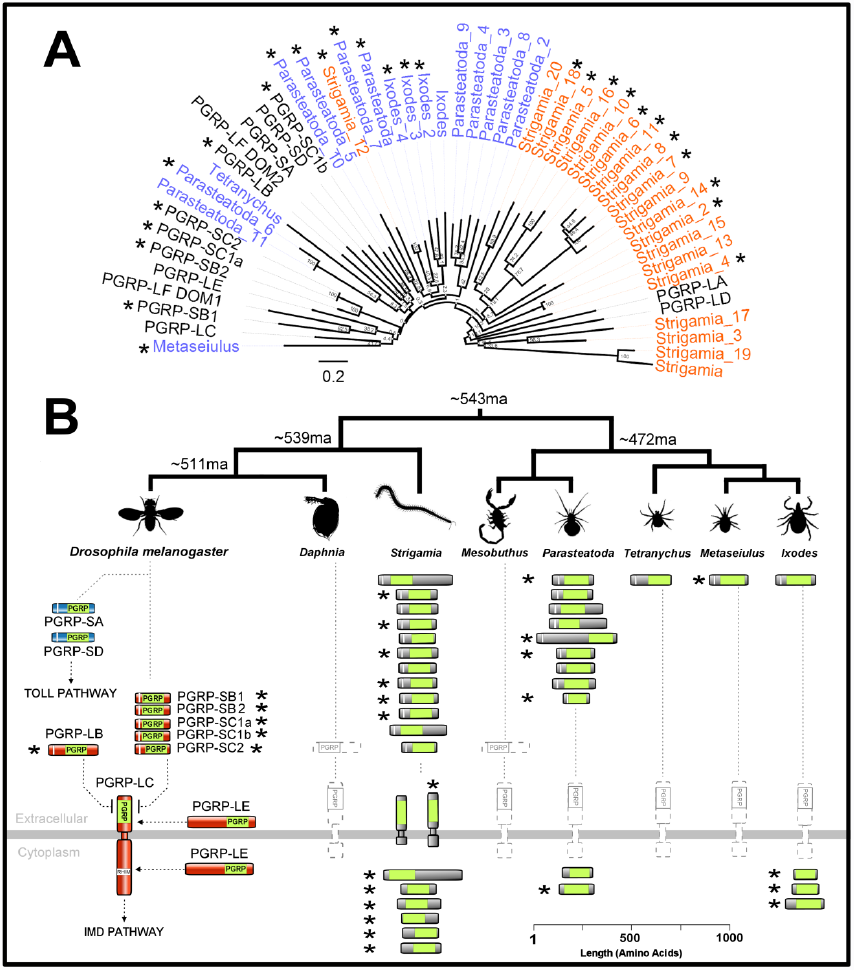
Gene tree, copy number, cellular location and predicted catalytic activity of arthropod PGRPs. (A) PGRP tree was reconstructed by maximum likelihood tree from the PGRP domain sequences and is midpoint rooted. Myriapod taxon labels are orange, chelicerates blue and *Drosophila* black. Node labels are bootstrap support. (B) Scale drawing and predicted cellular location of PGRPs. The PGRP domain is shown in lime green, signal peptides in white. Predicted catalytic PGRPs are denoted by an *.

The copy number of PGRPs varies greatly across the arthropods (Figure 4). They have been entirely lost from both the crustacean *Daphnia* (see also(McTaggart et al. 2009)) and the chelicerate *Mesobuthus*, while *Metaseiulus* and *Tetranychus* each have just a single PGRP. At the other extreme the myriapod *Strigamia* has 20 PGRPs. All bar one of the *Strigamia* genes cluster together on the gene tree, which suggests that they have probably resulted from one or two ancestral PGRPs that duplicated extensively within the myriapods (Figure 4).

It is possible to predict whether a PGRP has catalytic activity from the sequence of its PGRP domain. The catalytic (amidase) activity of insect PGRPs and bacteriophage T7 lysozyme is zinc-dependant, and the non-catalytic PGRPs have lost residues required for zinc binding(Mellroth, Karlsson, and Steiner 2003; Reiser, Teyton, and Wilson 2004; Gendrin et al. 2013).To predict the function of the PGRPs, we aligned their PGRP domains and identified a cysteine and two histidine residues (see Figure 3 of (Reiser, Teyton, and Wilson 2004)) that are required for zinc binding(Mellroth, Karlsson, and Steiner 2003; Reiser, Teyton, and Wilson 2004). In support of the functional relationship between these sites, we found that the presence of these three residues was strongly correlated – 18 of the 21 sequences with the cysteine also had the two histidines, while only one of the 16 sequences without the cysteine had both histidines (Supplementary Table 2).

Catalytic and non-catalytic PGRPs are scattered across the arthropods, and every species with more than one PGRP has both types (Figure 4, catalytic forms marked *). There have been frequent gains or losses of catalytic activity during evolution, as the two forms are interspersed on the gene tree, and closely related pairs of sequences can differ in whether they have predicted catalytic activity (Figure 4). Together these results suggest that in other arthropods, as in *Drosophila,* PGRPs are likely to play a variety of roles. However, as catalytic or non-catalytic PGRPs are absent from taxa across the tree, it seems likely that these functions may be replaced by other molecules in these groups.

In chelicerates and myriapods many of the PGRPs do not have a signal peptide and are therefore predicted to be intracellular (Figure 4). In *Drosophila,* intracellular isoforms of PGRP-LE are important in responses to intracellular bacteria(Kaneko et al. 2006), and can both activate the Imd pathway and induce autophagy in an Imd-independent way(Yano et al. 2008). It is unlikely the intracellular PGRPs that we identified perform similar roles, as they differ from PGRP-LE in that they are predicted to be catalytic and do not contain a predicted RHIM motif that could interact with Imd. Therefore, these intracellular catalytic PGRPs may have a novel function such as killing intracellular bacteria.

### Beta 1,3 Glucan Recognition Proteins (BGRP) have been lost from chelicerates

Beta-1,3 Glucan Recognition Proteins (BGRPs), which are also known as Gram negative binding proteins (GNBPs), bind Beta-1,3 glucan in microbial cell walls (particularly fungi), and can act as co-receptors with PGRP-SA and SD to recognise Gram-positive bacteria and initiate the *Drosophila* Toll pathway through Spatzle. Unlike the widely taxonomically distributed PGRPs, we find proteins bearing the functionally diagnostic GH16-superfamily domain to be limited to the Mandibulata (insects, crustaceans and myriapods) and entirely absent from any of the chelicerates (Supplementary figure 1). This pattern suggests a single loss event on the branch leading to chelicerata, as BGRPs are known to be present in molluscs, an outgroup to both these subphyla (Zhang et al. 2012). *Drosophila* and *Strigamia* both possess three GH16-bearing proteins, while *Daphnia* has ten. From phylogenetic analysis it would appear that these are the result of a lineage specific expansion in crustaceans (Supplementary figure 1).

### Arthropod thioester-containing proteins include relatives of vertebrate C3 complement factors and proteins lacking the thioester motif

The TEPs include the vertebrate complement factors C3, C4 and C5, the insect TEPs, and a family of vertebrate protease inhibitors called alpha-2-macroglobulins. We found members of the thioester-containing protein family (TEPs) in all species, confirming that they were present in the common ancestor of the arthropods and have been retained in all the major arthropod lineages (Figure 5).

In *Drosophila* and mosquitoes, TEPs can covalently bind to the surface of pathogens and parasites, and mark them for destruction by phagocytosis or melanotic encapsulation(Levashina et al. 2001; Blandin et al. 2004; Stroschein-Stevenson et al. 2006). The TEPs have a characteristic thioester motif, which once the TEP has been cleaved into its active form can covalently bind to pathogens(Blandin et al. 2004; Sekiguchi, Fujito, and Nonaka 2012). We found TEPs bearing the thioester motif in all species (Figure 5).

All the arthropod genomes also encode TEPs that lack the canonical thioester motif GCGEQ, and therefore presumably lack the ability to form covalent thioester bonds to microbial surfaces (Figure 5, supplementary table 5). Importantly, most of these proteins lack the critical cysteine required for the formation of thioester bonds (supplementary table 5) All but two of these fall into a single clade, all the members of which lack this motif (Figure 5, highlighted green). This clade includes the *Drosophila* protein MCR (macroglobulin complement related or Tep VI), so we have named these macroglobulin complement related (MCR) proteins.

**Figure 5.**
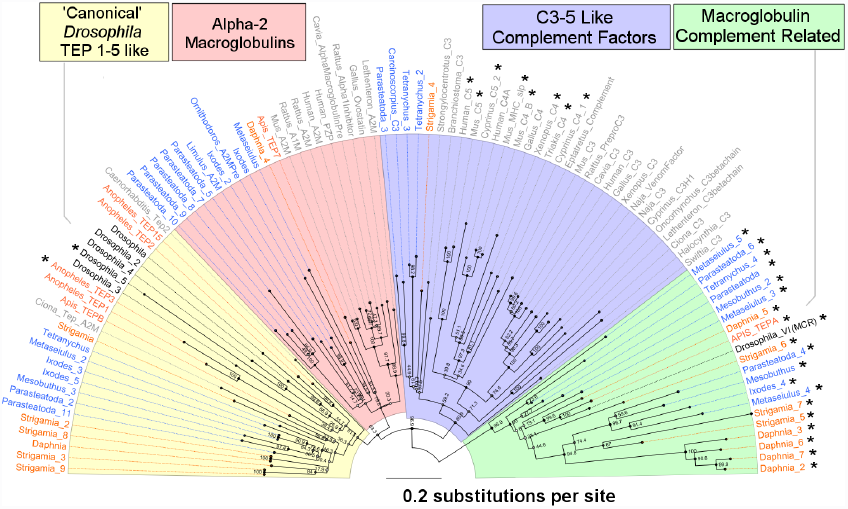
Gene tree of the thioester-containing protein (TEP) family. Sequences include arthropod TEPs, macroglobulin complement related proteins (MCRs), the vertebrate C3, C4 and C5 complement factors, and alpha-2 macroglobulins. Genes without a thioester motif are shown with an ‘*’. Myriapod and crustacean taxon labels are orange, chelicerates blue and *Drosophila* black. Taxa in grey are all deuterostomes with the exception of the nematode *Caenorhabditis*, which is a protostome related to arthropods, and the coral *Swiftia*, which diverged before the split of vertebrates and arthropods. In addition to the chelicerate sequences we annotated, we included two arthropod sequences from horseshoe crabs (*Limulus* and *Carcinoscorpius*) and a sequence from the tick *Ornithodoros.* The tree is midpoint rooted and was reconstructed by maximum likelihood. Node labels are bootstrap support. The additional taxa are taken from the previous analyses of Zhu et al, 2005 (Zhu et al. 2005); Wu et al, 2012; and Sekiguchi et al, 2012 (Sekiguchi, Fujito, and Nonaka 2012).

Despite lacking the thioester motif, in *Drosophila* MCR can bind to the fungus *Candida albicans* and promote phagocytosis(Stroschein-Stevenson et al. 2006). Our results show that MCR proteins were present in the common ancestor of arthropods and all eight of the species that we studied have at least one copy, with *Daphnia* having four copies and *Strigamia* and *Parasteatoda* having three copies (Figure 5).

Phylogenetic analysis revealed proteins related to vertebrate C3 complement factors in the myriapod *Strigamia* and the chelicerates *Tetranychus* and *Parasteatoda* (Figure 5). The complement factors fall into a monophyletic group containing a single clade of vertebrate complement factors, a clade of arthropod sequences, and a basal lineage found in corals (Figure 5). The relationships of these clades therefore mirror the phylogeny of these three groups, and indicate an ancient origin of the C3 complement factors. These results support a previous finding of a sequence from the chelicerate *Carcinoscorpius* that was most similar to C3 (ref^5^; Figure 5). Therefore C3-like proteins are widely scattered across the arthropods, but have been lost from most lineages including *Drosophila*.

The remaining two clades include the vertebrate alpha2 macroglobulins and the *Drosophila* TEPs respectively (Figure 5, pink and yellow). Members of the *Drosophila* TEP clade are found in all the arthropod genomes we analysed, although species other than *Drosophila* only have one or two copies (Figure 5, yellow). The function of these proteins has been best studied in *Anopheles* mosquitoes where they can bind bacteria and eukaryotic parasites, promoting phagocytosis and encapsulation(Levashina et al. 2001; Blandin et al. 2004). Interestingly *C. elegans Tep2* is a sister to this monophyletic group of arthropod TEPs. Sister to this group of insect TEPs we found a clear monophyletic grouping of vertebrate alpha-2 macroglobulins and arthropod sequences (Figure 5, pink). Of our sequences, we found five *Parasteatoda*, two *Ixodes*, one *Metaseiulus* and one *Daphnia* sequence to be alpha-2 macroglobulin-like. An insect sequence — TEP7 annotated in the honey bee genome project(Evans et al. 2006)— also fell into this clade.

### Gene duplication generates diversity in the immune receptor Dscam

Dscam (Down Syndrome Cell Adhesion Molecule) can bind to bacteria and promote their phagocytosis in *Drosophila*, and it is remarkable in that alternative splicing can potentially produce in the order of 38,000 isoforms(Watson et al. 2005). Such extreme receptor diversity was previously thought to be restricted to the vertebrate immune system, although it is unclear to what extent it is required for Dscam’s immune function or for its role in neuronal development. The Dscam homolog in *Daphnia* also generates diversity though alternative splicing(Brites et al. 2008), while a recent study found that *Strigamia* has the potential to express a high diversity of Dscam receptors through extensive paralogous gene duplication combined with a lower level of alternative splicing(Brites et al. 2013).

There are multiple Dscam copies in all the species we studied, and frequent duplications or losses of the gene (Figure 6). Lineage specific duplications of Dscam have resulted in 60 copies in *Strigamia* (a similar number were previously reported in this species(Brites et al. 2013)) and 35 in the spider *Parasteatoda*, while in the other species we find four to fourteen copies (Figure 6). While we did not characterise the diversity generated through alternative splicing, these results indicate that Dscam diversity is an important trait that is generated in different ways across the arthropods.

**Figure 6.**
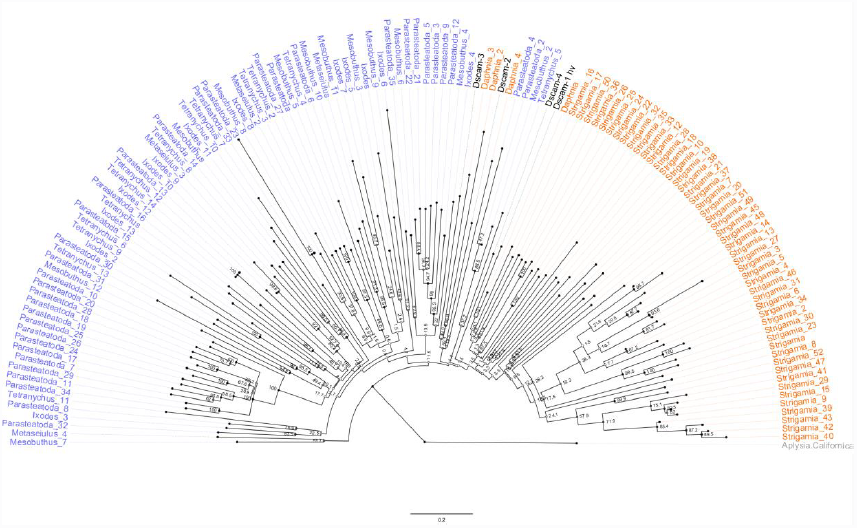
The diversity of Dscam in the arthropods. The tree is reconstructed by maximum likelihood based on an amino acid sequence alignment and is rooted using the mollusc *Aplysia californica*. Myriapod and crustacean taxon labels are orange, chelicerates blue, and *Drosophila* black. Node labels are bootstrap support.

### Fibrinogen-related proteins (FREPs) and Nimrod-like proteins

All species had FREPs, which are a family of proteins that contain a conserved fibrinogen domain and are involved in many anti-pathogen processes, including binding to pathogens and acting as pattern recognition receptors in phagocytosis and vertebrate complement activation(Dong and Dimopoulos 2009). The copy number of FREPs is highly variable, with the mites *Metaseiulus* and *Tetranychus* containing one and two fibrinogen domain containing proteins respectively, whilst all other genomes contain between nineteen and thirty-six (table 3).

Genes of the Nimrod superfamily are characterised by the presence of NIM-repeats, a specialised type of the EGF-domain(Somogyi et al. 2008). They have previously been shown to be widely taxonomically distributed and are known to have roles in phagocytosis in insects, nematodes and humans(Somogyi et al. 2008). We found multiple Draper like proteins in all genomes except *Metaseiulus* and *Mesobuthus*, which likely represent secondary losses or a lack of detection power. We failed to identify any Nimrod B-type or C-type (eg *Eater in Drosophila*) proteins in any group beyond *Drosophila*.

### Prophenoloxidase, nitric oxide synthase and DUOX

Dual oxidase (DUOX), which can produce reactive oxygen species that kill microbes in *Drosophila*(Ha et al. 2005) was present in variable copy number in all taxa (Supplementary table 3). Nitric oxide synthase (NOS) is an important enzyme in *Drosophila* with diverse roles in development and immunity. In insects, nitric oxide and NOS have been shown to be involved in both the direct killing of parasites and immune signalling(Foley and O’Farrell 2003). We find homologues of *Drosophila* NOS in all taxa except *Metaseiulus*.

Melanin is essential in the encapsulation of parasites, wound healing and the hardening of cuticle, and is produced when the inactive zymogen prophenoloxidase (PPO) is cleaved into its active form by a serine protease cascade, resulting in the oxidation of phenols. The three *D. melanogaster* PPOs each bear Hemocyanin N, Hemocyanin M, and Hemocyanin C domains in that order. Hemocyanin M additionally has a tyrosinase motif, which may distinguish it from closely related proteins such as hexamarins and larval storage proteins(Burmester 2001). We find predicted proteins bearing the three syntenic domains in *Daphnia, Strigamia, Mesobuthus,* and *Parasteatoda* (although conserved domain searches only predicted tyrosinase activity in the *Parasteatoda* and *Mesobuthus* homologues; Table S3). This protein family was greatly expanded in the arachnids, with 13 copies in *Parasteatoda* and 8 in *Mesobuthus*. These proteins may be functioning as PPOs or oxygen-carrying hemocyanins.

## Conclusions

Our results confirm an ancient origin for the innate immune system, predating the split between protostomes and deuterostomes. We find striking examples of conservation between vertebrates and arthropods, despite these two groups having diverged before the Cambrian explosion some 543 million years ago. These include a group of arthropod toll-like receptors that share structural similarity with vertebrate TLRs and cluster with them phylogenetically. Similarly, several arthropods have C3-complement like factors that have been lost from *Drosophila*.

Despite such conservation, we also find remarkable diversity in the immune system of different arthropods. The Imd pathway - essential for recognition and response to gram negative bacteria in *Drosophila* - is almost entirely absent from the chelicerates. We also observe extensive copy number variation in recognition and effector genes, suggesting complex evolutionary dynamics in these functional categories. Detailed analysis of PGRPs suggests that this change in gene copy number is accompanied by changes in function.

This is the first detailed genome-wide analysis of arthropod immune systems outside of insects. The non-insect arthropods comprise a significant proportion of the earth’s biodiversity and include many species of economic and medical importance. Characterising the function of the immune genes that we described remains an important challenge for the future.

## Methods

### Data and Query set

We retrieved the complete predicted-peptide sets for *Drosophila melanogaster* (r5.54, Flybase) and six additional non-insect arthropods: *Strigamia maritima* (v1.20, Ensembl Genomes), *Metaseiulus occidentalis* (v1.0, NCBI Refseq), *Tetranychus urticae* (v1.2, Ensembl Genomes,(Grbić et al. 2011), *Ixodes scapularis* (v1.2, Vectorbase), *Daphnia pulex* (r20, Ensembl Genomes,(Colbourne et al. 2011)), *Parasteatoda tepidariorum* (Augustus 3, Spiderweb) and performed all analyses on these data. A query set of key immunity genes in *D. melanogaster* was compiled from Immunodb (Waterhouse et al. 2007), IIID (Brucker et al. 2012), Flybase (Pierre et al. 2014) and (Obbard et al. 2009) (Supplementary Table 4).

### Identification of homologues

Broadly, to identify sequence homologues of a query set of *D. melanogaster* immunity genes (Supplementary table 4) we used multiple Blast-based and hidden Markov model based approaches, to compile a redundant list of candidate homologues in each of the six additional non-insect arthropod species. This list was then filtered by similarity, quality, e-value, best reciprocal *Drosophila* hit, presence/absence of conserved domains known to be essential to function, and additionally by tree-based similarity measures, producing a final non-redundant list of high confidence predicted *Drosophila* innate immunity functional homologues in each peptide set.

#### Ortholog clustering

To identify clusters of orthologous genes, we performed all-versus-all blast-based clu
stering of protein sequences using Orthomcl V1.4 (Li, Stoeckert, and Roos 2003) and the predicted peptide sets from all eight arthropod species. Default parameters were used, and homologues from each of the seven non-insect arthropods were assigned to immune homology groups based on clustering with a *Drosophila* innate immunity peptide from our query set.

### HMMER

For a subset of genes we searched for homologs in each full peptide-set using the hidden Markov model based HMMER (Eddy 2011). HMMER aims to be more sensitive than a traditional whole-sequence based search such as Blast, by utilising profile HMMs. Profile HMMs are statistical models of multiple sequence alignments whereby each residue in the alignment is determined to be more or less relevant to homology based on its conservation between sequences. Hits with an e-value of less than 10^-5^ were considered significant and retained.

We built profile HMMs for each of the eighteen signalling pathway peptides (Toll, Imd, JNK, and JAK/STAT), using multiple alignments of the *Drosophila melanogaster* gene and its high-confidence orthologues in the immune-annotated insects *Bombyx mori, Tribolium castaneum,* and *Apis melifera* (Tanaka et al. 2008; Zou et al. 2007; Evans et al. 2006). Multiple alignments were built using MAFFT(Katoh, Asimenos, and Toh 2009), and orthologous sequences were retrieved from Tanaka et al, 2008 (*B. mori*), Zou et al, 2007 (*T. castaneum*) and Evans et al 2006 (*A. melifera*).

For the highly variable and divergent Nimrod genes (for which blast/orthomcl does not find significant homologues, and no single diagnostic domain exists), we built a profile HMM of the conserved NIM motif CXPXCXXXCXNGXCXXPXXCXCXXGY(Kurucz et al. 2007; Somogyi et al. 2008), using the multiple alignment of (Somogyi et al. 2008) and searched the complete peptide sets using HMMER (Eddy 2011).

#### Blastp

We also searched all seven non-*Drosophila* peptide-sets for homologues of *Drosophila* immune genes using Blastp, retaining hits with an e-value less than 10^-6^, greater than 20% identity, and a bit score > 80. We additionally queried each species’ complete peptide-set against a blast database of all *D. melanogaster* peptides, in order to identify the best *Drosophila* blastp hit for each gene in each other species.

#### Identification of protein domains

Some gene classes rely on a conserved protein domain to function in the manner described in *D. melanogaster,* without which the protein was assumed to be non-functional and therefore discarded. We searched each whole non-*Drosophila* peptide-set for domains known to function in innate immune pathways using the NCBI BatchConservedDomain tool and identified proteins containing domains listed in the query set (Supplementary table 1).

## Combining results and further analyses

We finally compiled results from all the above techniques into a single list of potential immunity homologues. Examining these results individually, we filtered out hits where either an essential protein domain was absent (Supplementary table 1) or where the reciprocal top *Drosophila melanogaster* blast hit was to a non-immunity related gene.

For PGRPs, GNBPs/BGRPs, and TLRs we scanned putative homologues for transmembrane helices and signal peptides using the TMHMM Server v. 2.0 (Krogh et al. 2001) and the SignalP 4.1 Server (Petersen et al. 2011) respectively. We also built a profile HMM of the PGRP RHIM (Imd-binding) domain after (Meister et al. 2009), and scanned putative PGRPs in all species to predict presence/absence of the RHIM domain.

Toll like proteins/receptors are characterised by an N-Terminal TIR domain, a transmembrane helix, and a variable number of leucine rich repeats (LRRs) extending to the C-terminal. TIR domains and transmembrane helices were identified as above, whilst LRRs were identified using the web-interface of LRR-finder (Offord, Coffey, and Werling 2010).

We additionally built trees of gene families where duplications were suspected to be assembly errors (eg. haplotypes annotated as separate contigs), or to distinguish orthologues and paralogues. Multiple alignments were assembled using MAFFT(Katoh, Asimenos, and Toh 2009), with phylogenetic tree construction performed in PHYML (Guindon et al. 2010) using the WAG+G model. Shimodaira-hasegawa tests to compare the log-likelihood of differing tree topologies were performed in RAxML (Stamatakis 2014) using the WAG+G model and multiple alignments created as above in MAAFT.

## Acknowledgements

This project was funded by a Royal Society University Research Fellowship and a European Research Council grant DrosophilaInfection to FMJ, and a Medical Research Council studentship to WJP. We also thank Marjorie Hoy and Alistair McGregor for early access to the *Metaseiulus* and *Parasteatoda* protein sets respectively.

